# Enhancing Sustainable Agriculture in Sub-Saharan Africa: The Potentials of Entomopathogenic Nematodes for Pest Control

**DOI:** 10.1101/2023.12.30.573611

**Authors:** Osawe Nosa

## Abstract

Entomopathogenic nematodes (EPNs) are a type of soil-dwelling nematodes with a remarkable ability to infect insect pests. Their employment for control of pests is a means to foster sustainable agricultural practices. These environmentally safe alternatives to pesticides are being researched in Sub-Saharan Africa (SSA), but there are many gaps to be filled, which this review exposes. The systematic analysis of peer-reviewed articles from 2003 to 2022 shows that about 56% of published articles on EPNs in Sub-Saharan Africa (SSA) come from studies in South Africa. In SSA, applied pest management studies of these nematodes have focused on Lepidopteran (41.7%), Coleopteran (22.2%), and Hemipteran (18.1%) pests, with *Cydia* sp., *Planococcus* sp., and *Thaumatotibia* sp. being the most targeted genera. Many areas in SSA remain unexplored for EPNs and knowledge of their use is still experimental. There is a paucity of knowledge regarding the bacteria symbionts of indigenous EPNs in SSA, with the possibility of variations in bacterial symbionts being associated with specific ecological populations. However, promising strides have been made, and further surveys are needed to explore untapped geographies. Utilizing native EPN species holds promise in curbing the reliance on hazardous pesticides and promoting integrated pest management approaches in SSA, representing a sustainable method for pest control.

## MAIN BODY INTRODUCTION

Entomopathogenic nematodes (EPNs) are small, soil-dwelling roundworms that are parasitic to insects; their infective juvenile (IJ) stage can kill and replicate within the host body. These potentials have been harnessed for biologically controlling insect pests in agriculture, horticulture, and forestry sectors (Farrar et al., 2016). They have effectively managed pests of important crops such as maize, palm, and rice (Mwaniki et al., 2008; Rosa et al., 2000). Also, their compatibility with other biocontrol agents and their safety in application substantiates their importance for integrated pest management (Abate et al., 2017; Georgis et al., 2006; Mutegi et al., 2017b).

To achieve sustainable agriculture, pest management should target reducing economic, health, and environmental risks. There has been a subsequent increase in the use of EPNs for the management of pests worldwide due to their eco-friendliness (Abate et al., 2017). EPNs’ ease of application through conventional liquid application equipment; such as those used in pesticide application make it a suitable pest control agent for farmers (Abate et al., 2017).

Despite the many potentials of EPNs, there is limited knowledge about the genetic and ecological diversity, and thus, the potentials of local EPNs in SSA (Grewal et al., 2001; G. Houssou et al., 2014). Scientific efforts to understand EPN distribution vary across regions. While substantial surveys and isolation have been made in some countries, and some have begun commercialization, many more have not reported any indigenous EPN (Abate et al., 2017). The potential of local EPNs would be harnessed when there is enough understanding to improve their infectivity and virulence to target insects. In this review, the level of knowledge of the distribution of species of EPNs in SSA is discussed by emphasizing what we know and suggesting areas where more research is needed. This review will also specifically cover the significance of the environmental characterization of local EPNs, the necessity of taxonomic and molecular characterization, and the possibility of utilizing EPNs as a sustainable alternative to chemical pesticides.

## MATERIALS AND METHODS

To extract relevant articles, Google Scholar and Scopus databases were utilized by employing a targeted search strategy based on keywords related to entomopathogenic nematodes. Specifically, this set of keywords were: ‘Entomopathogenic ‘XXX’ nematode OR nematodes OR *Steinernema* OR *Heterorhabditis* OR *Oscheius*’. Here, “XXX” represents the name of a specific country in SSA. To ensure that only peer-reviewed articles were included, the search results were thoroughly screened by reviewing the title, abstract, and materials and methods sections. Only articles which focused on research related to local EPNs and/or the use of EPNs for the management of local pests were reviewed. Furthermore, only papers published within the years 2003-2022 were included. If 20 consecutive search results did not meet the inclusion criteria, it was assumed that the remaining search results would be irrelevant and stopped screening further. Finally, duplicate articles were excluded from the review.

## RESULTS

The analysis of 151 peer-reviewed published literature for EPN in SSA (which passed the inclusion criteria) shows that the majority of the locally isolated EPNs have been identified within the past decade (Fig 1). However, going by the described nematodes so far, the native EPNs in SSA belong to one of three genera: *Heterorhabditis*, *Oscheius*, and *Steinernema*, with *Oscheius* spp. being more recently reported in the region (Daramola et al., 2022; Lephoto & Gray, 2019). Notably, the majority of pest control studies have targeted Lepidopteran (41.7%), Coleopteran (22.2%), and Hemipteran (18.1%) insects (Fig 2). Also, the publication records show *Cydia* sp., *Planococcus* sp., and *Thaumatotibia* sp. are the insect pests most frequently targeted by EPNs for management purposes (Fig 3).

**Figure 1:**
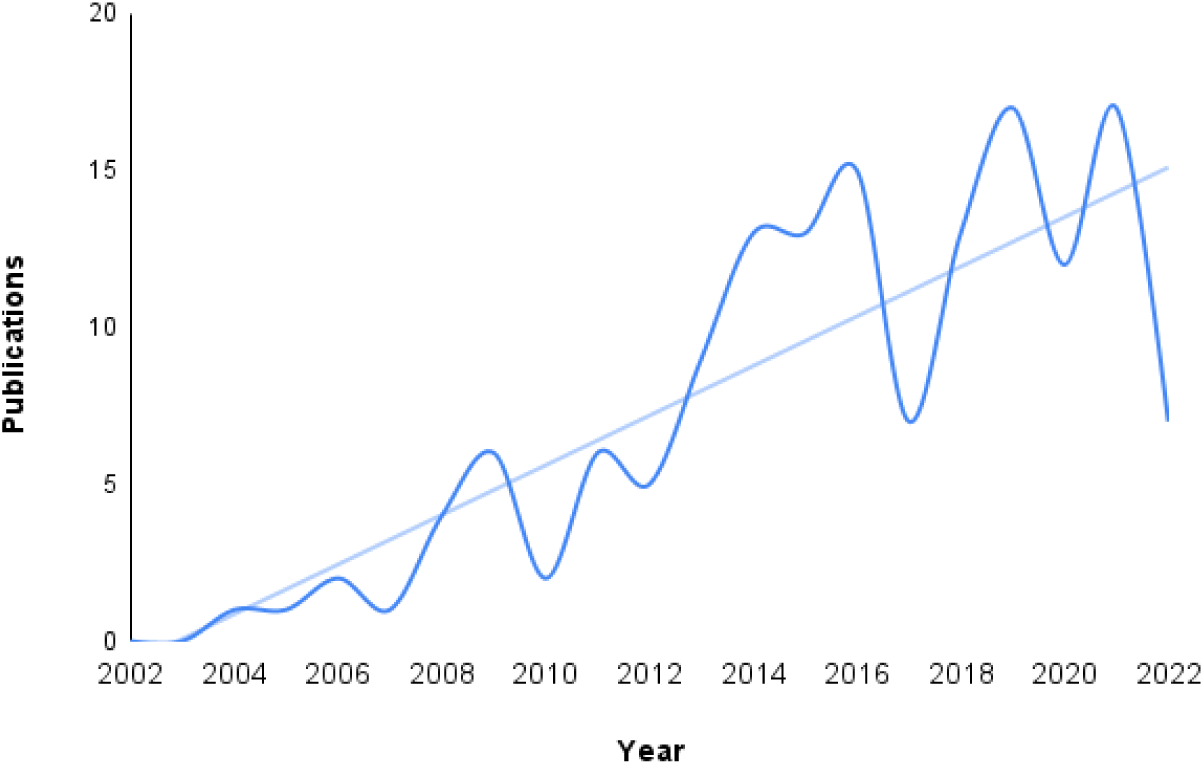
Number of yearly peer-reviewed EPN-related publications from SSA

**Figure 2:**
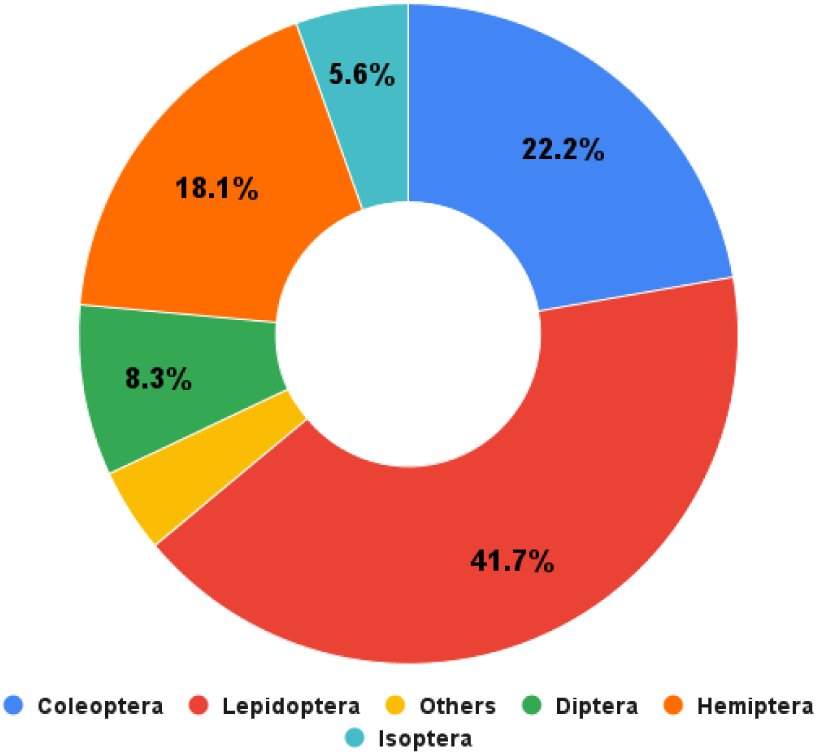
Pest management focus in EPN publications for SSA (2003-2022)

**Figure 3:**
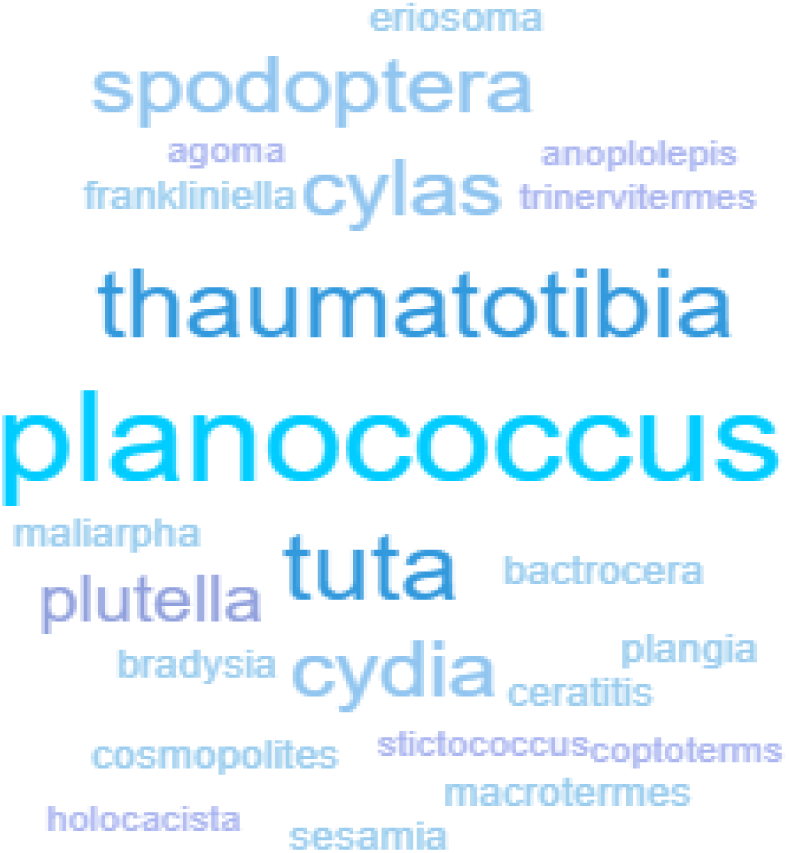
Word cloud showing target pest genera in EPN publications from SSA countries (2003- 2022)

The results show that the investigations on EPNs in SSA are expanding, although it is at a preliminary level (Fig 1). However, the bulk of our understanding of local EPNs in SSA arises from investigations on South African EPN isolates, accounting for 56% of published articles (Fig 4). Moreover, there has been a remarkable increase (Fig 1) in our knowledge of EPNs in SSA, with newer EPN species described in the past decade than ever before. In most West African countries, the first description of their EPN isolates at the species level occurred after 2010. In 2015, Aliyu reported the first isolation of a nematode-bacteria complex in West Africa (Aliyu, 2015). Houssou (2012) also published the first investigation into the characterization of the bacterial complex of EPNs in SSA, which was conducted in Benin. The first detection of EPNs in Nigeria was reported in 2012, including *H. bacteriophora* and *S. feltiae* isolates (Akyazi et al., 2013). Daramola et al. (2021) made the first report on the molecular characterization of *Heterorhabditis* sp. in Nigeria, providing valuable insights into the genetic diversity of EPNs in the region.

**Figure 4:**
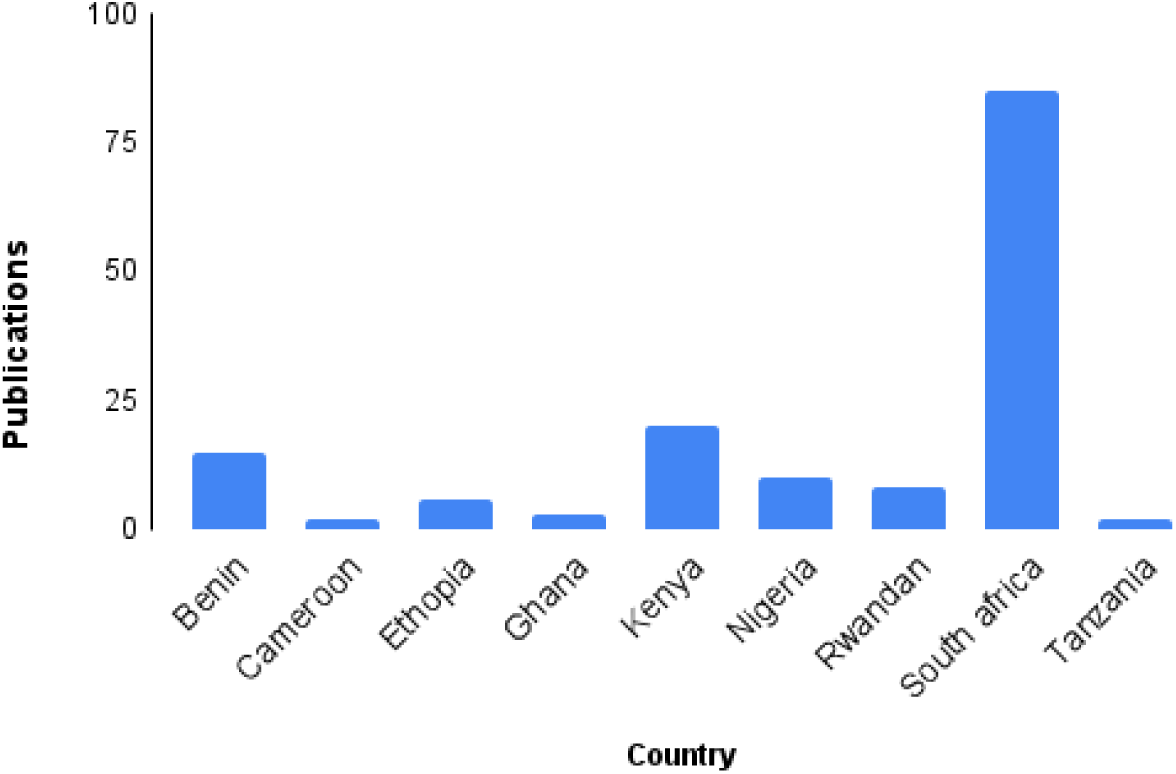
Number of peer-reviewed publications for SSA countries from the year 2003-2022

Although there is a paucity of knowledge on the indigenous EPNs in SSA, research publications on these insect-killing nematodes in the region have increased (Fig. 1). Results show that EPN isolation has been carried out in several SSA countries, including Kenya, Ghana, Ethiopia, Tanzania, Benin, South Africa, Rwanda, Cameroon, and Nigeria (Fig. 4). However, only a handful of countries in the region, including South Africa, Kenya, Benin, and Nigeria, have published EPN surveys from more than five geographic locations.

### Potential Benefits of EPN in SSA

The employment of EPN in SSA for pest management would foster sustainable agriculture and biodiversity (Akyazi et al., 2013; Divya & Sankar, 2009). These nematodes are useful for pest control and resistance management initiatives due to their minimal environmental effect and selectivity (Georgis et al., 2006; Mutegi et al., 2017a). The nematode-bacterium complex can kill pests within short periods and so does not give enough time for the establishment of host-parasite relationships. This ‘kill-quick’ action does not provide room for the host to develop defences against the parasite, unlike most other nematode associations (Divya & Sankar, 2009; Peters, 1996). Some EPN infective juveniles have high persistency in the ecological environment and are of global reuse due to their ability to multiply and emerge from a dead host (insect) (Baimey et al., 2017; Fanou et al., 2019; Godjo et al., 2018). Furthermore, EPNs are more effective than most microbial control agents at controlling various pests (Divya & Sankar, 2009; Peters, 1996).

Many EPNs can be integrated with biopesticides for effective pest control (Laznik & Trdan, 2014; Özdemir et al., 2020). For example, Mutegi *et al*. (2017) successfully integrated Neem (Azadirachtin 0.03%) with EPNs for the control of leaf miner (*Tuta Absoluta*); a pest of tomatoes. It is noteworthy that compatibility is both species-specific and sometimes strain-specific; and understanding of the species should be, before integration with pesticides by growers for pest management (Claudius-Cole, 2018; Divya & Sankar, 2009; Georgis et al., 2006).

Farmers acquainted with conventional pest management practices may not need special training on the use of equipment for EPN application in their farms. EPNs will benefit growers in areas where pesticides are not accessible and/or too toxic or not practical. As a consequence, consumers will also benefit from foods with less pesticide residue (Yan et al., 2016).

Numerous accounts of the successful utilization of EPNs to control indigenous pests in locations within SSA have been documented. Morris et al. (2020) investigated the susceptibility of *Agoma trimenii* (Lepidoptera: Agaristidae) larvae and pupae to two South African isolated EPNs, *S. yirgalemense* and *H. noenieputensis*. They added the infective juveniles (IJ) to water at a concentration of 100IJs/50ul of water which was used for the treatment of the pest (*A. trimenii*), for which they observed 100% mortality when applied to the larvae stage. Red Palm Weevil (RPW), *Rhynchophorus ferrugineus* (Coleoptera: Curculionidae), is obnoxious to stem tissues of oil palm trees in West Africa. In Nigeria, Ottun et al. (2021) inoculated larvae of *R. ferrugineus* with native *Heterorhabitis* sp. and observed that 84% of the pest died after 9 days post-inoculation. A study carried out at Kenya’s Agricultural Research Station, Mwea, showed that local EPNs were efficient in the management of African rice borer (*Maliarpha separatella*) larvae. Kega et al. (2020) showed that *H. indica* (at concentrations up to 200 IJ/larva) was highly effective against *M. separatella* within 48 hours of infection. Also, the Diamond black moth (DBM) is a crop pest that causes extreme yield losses and is conventionally controlled using pesticides in SSA. Nyasani et al. (2008) found that foliar application of *H. indica, S. weiseri,* and *Steinernema* sp. effectively killed DBM larvae 48 hours after inoculation in field experiments. Odendaal et al. (2016) conducted experiments in South Africa to evaluate the efficacy of imported and local *S. feltiae*, *H. bacteriophora*, and *S. yirgalemense* isolates in managing codling moth under local conditions. The infectivity of both imported *S. feltiae* and local *S. yirgalemense* isolates was high when applied at a concentration of 1000 IJ/ml, 24 hours post-application. In addition, the recycled imported *S. feltiae* showed significantly higher virulence than the local *S. yirgalemense* isolates under the same exposure conditions.

### Challenges with the implementation of EPN for pest control in SSA

Local climatic conditions, specifically moisture and humidity levels, have a significant impact on the effectiveness of EPNs in pest control. Employing endemic EPN is recommended due to their adaptation to prevailing climatic conditions, making them better suited for controlling the local pest population. This approach promotes sustainable and environmentally friendly pest management (Odendaal et al., 2016; Yan et al., 2016). Environmental characterization of local EPNs is necessary for selecting the best formulation design for pest control. EPNs typically thrive at temperatures below 30°C, but they are intolerant at temperatures higher than 35°C (Grewal et al., 1994; Odendaal et al., 2016; Platt et al., 2020). Thus, formulations that protect nematodes from environmental stress should be researched to increase the efficiency of EPNs for biocontrol (Grewal, 2002). The relative humidity should be high, and the ambient temperature should be neither extremely hot nor cold, with soil temperature between 10-35°C, moist soil, and minimal direct sunlight to prevent nematode death and enhance their survival and virulence (Grewal et al., 1994; Odendaal et al., 2016; Platt et al., 2020).

The most prevalent reason for failure in EPN applications has been a poor understanding of the nematode and target pests’ interaction (Gaugler, 1999; Georgis et al., 2006). Biological characteristics such as resistance to desiccation and temperature tolerance are also crucial in selecting the best-adapted EPN pests (Georgis et al., 2006; Yan et al., 2016). In most cases, the pest’s larval mortality observed in the field is lower than that seen in laboratory conditions (Nyasani et al., 2008). The pathogenicity of EPNs is also dependent on soil moisture content and aeration (Mwaniki et al., 2008; Odendaal et al., 2016; Rufai et al., 2020). Furthermore, being soil-dwelling they are less effective in managing above-ground pests due to desiccation of the free-living infective juveniles. Also, there are concerns about the possible non-target effects of EPNs. Some argue that EPNs may harm hymenopteran parasitoids or beneficial predators in the field (Akhurst & Smith, 2002), and their use in pest management is hindered by an unfavourable climate (Platt et al., 2020). In SSA, the current understanding of local EPNs is largely based on laboratory experiments, which may not provide a complete representation of their field performance. There is a notable paucity of knowledge regarding the bacteria symbionts of indigenous EPNs in SSA, with the possibility of variations in bacterial symbionts being associated with specific ecological populations (Abate et al., 2017; Baimey et al., 2017).

EPN research in SSA is still in the experimental phase and funding is needed to light the practice of biological control with EPNs. However, South Africa is doing relatively well. The majority of published research done in SSA is from South Africa. This review shows that South Africa’s National Research Foundation and the Gauteng Department of Agricultural and Rural Development are major sponsors of its EPN research. The low level of research on EPNs in SSA is largely attributed to a lack of funding. The scarcity of professional experience in the subject has caused little attention to these nematodes as insect pest biocontrol agents in SSA. Funding EPN research in SSA would procure benefits such as; enhancement of knowledge, conservation biology, and creation of job opportunities.

### Commercialization of EPNs

The successful mass production of EPNs hinges on the selection of an appropriate artificial diet for rearing their host. The greater wax moth *Galleria mellonella* L. (Lepidoptera: Pyralidae) is a commonly used bait for EPN mass production due to its susceptibility, size, short lifecycle, ease of rearing on artificial diets comprised of multiple ingredients, and ability to be reared at suitable temperatures, leading to high yield (Abate et al., 2017; Kotchofa & Baimey, 2019). Diet is a critical factor influencing *G. mellonella* development, with the lipid composition (which serves as the main source of energy for the host) varying largely due to the type of diet consumed (Andalo et al., 2011). The quality of *G. mellonella* and thus the reproductive potential and pathogenicity of EPNs depend largely on the diet fed to the host (Kotchofa & Baimey, 2019).

SSA may still be far from building notable commercial companies for EPN production due to several challenges, including the need for temperature regulation, uncertainty in performance, price considerations, and lack of knowledge on how to use EPNs efficiently, among others. Nematodes are known to survive well at temperatures below 35°C as high temperatures tend to negatively affect the nematode’s virulence and shelf life (Georgis et al., 2006; Odendaal et al., 2016). This means that there may be a need for a cooling system, such as refrigerators to keep the EPNs in the best conditions. An array of challenges faces the setting up of EPN commercial companies in SSA such as product price, uncertain performance, refrigerated needs for most formulations, usage of inferior nematode species, and a lack of particular knowledge on how to use EPNs’ efficiently (Abate et al., 2017; Divya & Sankar, 2009; Georgis et al., 2006; Kaya & Gaugler, 1993; Odendaal et al., 2016).

Promotion of the use of EPNs for pest control in SSA will be facilitated by a conducive socio-political environment, and SSA nations should follow the example of American and European nations, where progress in the commercialization of EPNs has been achieved through collaborations among universities, industries, and the government (Abate et al., 2017). Field efficacy is one of the required components for commercialization. Other important factors are cost, storage, delivery, handling, mixing, coverage, competition, compatibility with grower practices, and profit margins to manufacturers and distributors. Careful assessment and consideration of these factors are essential for product development and market penetration (Abate et al., 2017; Georgis et al., 2006; Lacey & Georgis, 2012).

Biological control agents are known to have a limited share in the market. In the African system, most farmers rely on synthetic pesticides for pest control (Abate et al., 2017; Divya & Sankar, 2009; Georgis et al., 2006). Despite increased publication trends in the region, lots of gaps in knowledge exist. The dispersal and behaviour of local EPNs in SSA are not generally known. Also, there has been no study to investigate the effect of long-term persistence of local EPNs in the region.

Genetic improvements for EPNs could lead to more stable formulations and wider use as biocontrol agents, resulting in cost savings by enabling effective control at lower rates than currently used (Gaugler, 1999; Georgis et al., 2006; Lacey & Georgis, 2012). However, there is limited effort to characterize the bacterial complex in EPNs, and identification has mostly been through morphological examination. Therefore, a more profound understanding of their biology, including ecology, behaviour, and genetics, is needed to comprehend their successes and failures as biopesticides (Georgis et al., 2006; Javal et al., 2019; Odendaal et al., 2016).

## RECOMMENDATIONS

It is imperative to isolate more African EPN strains to identify those that are cost-effective for mass production, which has an impact on commercialization (Divya & Sankar, 2009; Lacey & Georgis, 2012; Ngugi et al., 2021). This can be achieved through field and semi-field studies, which should be conducted before introducing the nematodes for pest management (Baimey et al., 2017; Georgis et al., 2006; Mutegi et al., 2017b). Future formulation improvements can increase the options for deployment against a wider range of targeted pests in SSA (Georgis et al., 2006). Instead of relying on imported commercial EPN pesticides, SSA should focus on understanding local EPN species and finding means to mass-rear them. Failure to test imported pesticides can have negative impacts on local EPNS and other beneficial biocontrol organisms (Abate et al., 2017; Lacey & Georgis, 2012). Also, future studies should concentrate on the EPNs’ environmental resilience in local settings to ensure their effectiveness as biocontrol agents (Nyasani et al., 2008). The identification of species and strains that can withstand harsh environmental conditions such as high temperatures, UV radiation, and dryness will broaden the scope of using entomopathogenic nematodes for biological pest control (Grewal et al., 2001).

The method of EPN isolation (which is primarily by the use of *G. mellonella*) is almost the same for all surveys in SSA. *G. mellonella* may not have been a good host for some of these nematodes (Mwaniki et al., 2008). As a consequence, some negative sites may have been positive all along but the nematodes remained non-infective to the bait (*G. mellonella*). The number of local EPNs isolated in surveys in SSA is likely an underrepresentation of the true ecological diversity. Dry periods or seasons, when EPNs are less active, can reduce their pathogenicity, which may contribute to the observed underrepresentation of the EPNs in some studies (Georgis et al., 2006; Godjo et al., 2018; Yan et al., 2016). Thus, repetitive sampling and the employment of multiple isolation methods for native EPNs should be adopted to understand their spatial and temporal distributions to avoid these drawbacks.

Despite progress in EPN studies in SSA, more surveys are needed in unexplored geographic areas (Bhat et al., 2020) to isolate new species, study their ecological footprint, survival, and abundance in different climates, and test their efficacy in pest management. These surveys should cover more regions, including plantations and forest areas, to identify native EPN species (Bhat et al., 2020) that are resilient to local environmental conditions and could provide significant resources from biodiversity, environmental, and economic standpoints (Abate et al., 2017).

Utilizing native EPN species would serve as an alternative to synthetic pesticides, ideally suited for local pest control (Aliyu, 2015; Bhat et al., 2020; Claudius-Cole, 2018; Yan et al., 2016). End-user training in the correct use of nematodes and their incorporation into pest control programs should be prioritized (Georgis et al., 2006). This might be achieved by hosting seminars on EPNs, promoting cooperative international initiatives involving specialists, and teaching African scientists, professionals, and students in academic institutions. National and international agreements should convey both parties’ mutual interests in terms of exchanging experiences, materials, and information, as well as respecting national and international regulations relating to biodiversity protection and the interchange of biological control agents (Divya & Sankar, 2009; Georgis et al., 2006). More specifically, the government should support research into local EPNs to gain a better understanding of their ecological characteristics and efficacy in pest management.

## CONCLUSION

The potential of EPNs as a pest biocontrol agent in SSA is yet to be fully explored. Despite South Africa’s leading role in research on this topic due to relatively high funding, other countries need to participate in surveys and the isolation and characterization of the nematode-bacterial complex. Utilizing native EPN species has the potential to significantly reduce our reliance on harmful chemical pesticides and promote a more holistic approach to pest management known as integrated pest management. However, the benefits of utilizing EPNs as biocontrol agents over traditional chemical pest control methods remain a topic of debate. From an economic perspective, the superiority of EPNs may not be immediately apparent as conventional pesticides are widely accessible and comparatively low-priced. Moreover, farmers in Africa, who have been reliant on chemical pesticides, may face challenges in adopting new pest management strategies. However, the acceptance of EPNs as pest control agents is contingent upon a host of factors, besides their efficacy, and comprehending the biology of EPNs is imperative for enhancing their virulence toward target insect pests.

EPNs have already shown great promise as effective biocontrol agents for local pests in SSA, and further exploration and utilization of this unharnessed potential could bring significant benefits to sustainable agriculture and biodiversity conservation in the region.

## ACKNOWLEDGMENT

Special thanks to Dr. Saheed Jimoh for his encouragement and constructive criticism of the methodology for database search.

