## Supplemental material for "Enhancing Sustainable Agriculture in Sub-Saharan Africa: The Potentials of Entomopathogenic Nematodes for Pest Control"

**Author: Osawe Nosa***

**Department of Animal and Environmental Biology, Faculty of Life Sciences, University of Benin, P.M.B 1154 Benin City -Nigeria**

**Table s1: Target insect pests for biological control using Entomopathogenic nematodes (EPNs) in Sub-saharan Africa (SSA) between year 2003-2022.**

| Genus | Species | Country | Region | Pest Order | Year | |
|---|---|---|---|---|---|---|
| Agoma | Trimenii | South Africa | Southern | Lepidoptera | 2020 | http://www.scielo.org.za/pdf/sajev/v41n2/06.pdf |
| Anoplolepis | tenella | Cameroon | Central | Hymenoptera | 2020 | https://doi.org/10.1016/j.biocontrol.2020.104321 |
| Bactrocera | dorsalis | Benin | Western | Diptera | 2018 | https://doi.org/10.1016/j.biocontrol.2017.10.009 |
| Bactrocera | dorsalis | Benin | Western | Diptera | 2021 | https://doi.org/10.1016/j.cropro.2021.105754 |
| Bradysia | impatiens | South Africa | Southern | Diptera | 2018 | https://journals.co.za/doi/epdf/10.4001/003.026.0337 |
| Bradysia | species | South Africa | Southern | Diptera | 2018 | https://journals.co.za/doi/abs/10.4001/003.026.0001 |

| | | | | | | |
|---|---|---|---|---|---|---|
| Cacosceles | newmannii | South Africa | Southern | Coleoptera | 2019 | https://doi.org/10.3390/insects10040117 |
| Ceratitis | capitata | South Africa | Southern | Diptera | 2017 | https://doi.org/10.1016/j.cropro.2017.11.008 |
| Ceratitis | capitata | South Africa | Southern | Diptera | 2008 | https://doi.org/10.1016/j.jip.2008.09.007 |
| Coptoterms | formosanus | Kenya | Eastern | Isoptera | 2017 | https://www.ajol.info/index.php/jagst/article/view/219191/206835 |
| Cosmopolites | sordidus | Tanzania | Eastern | Coleoptera | 2011 | https://doi.org/10.1017/S1742758411000294 |
| Cosmopolites | Sordidus | Kenya | Eastern | Coleoptera | 2016 | https://ideas.repec.org/a/aoj/agafsr/v3y2016i1p29-36id160.html |
| Cydia | pomonella | South Africa | Southern | Lepidoptera | 2016 | https://journals.co.za/doi/abs/10.4001/003.024.0061 |
| Cydia | pomonella | South Africa | Southern | Lepidoptera | 2015 | https://journals.co.za/doi/epdf/10.10520/EJC176580 |
| Cydia | pomonella | South Africa | Southern | Lepidoptera | 2015 | https://www.cambridge.org/core/journals/journal-of-helminthology/article/abs/entomopath [illegible] trials/CC61F883756FB5185E31E4B4FC5A4CB3## |
| Cydia | pomonella | South Africa | Southern | Lepidoptera | 2016 | https://doi.org/10.1080/09583157.2016.1217393 |
| Cydia | pomonella | South Africa | Southern | Lepidoptera | 2011 | https://doi.org/10.1080/09583157.2011.607922 |
| Cylas | puncticollis | Benin | Western | Coleoptera | 2017 | https://www.entomoljournal.com/archives/2017/vol5issue6/PartH/5-4-307-961.pdf |
| Cylas | puncticollis | Benin | Western | Coleoptera | 2019 | https://doi.org/10.4314/ijbcs.v13i1.36 |
| Cylas | puncticollis | Kenya | Eastern | Coleoptera | 2009 | https://www.cabdirect.org/cabdirect/abstract/20093232109 |
| Cylas | Puncticollis | Kenya | Eastern | Coleoptera | 2009 | https://www.kalro.org/www.eaafj.or.ke/index.php/path/article/view/390 |
| Cylas | puncticollis | Kenya | Eastern | Coleoptera | 2009 | http://erepository.uonbi.ac.ke/handle/11295/26308 |
| Eriosoma | lanigerum | South Africa | Southern | Hemiptera | 2016 | https://journals.co.za/doi/abs/10.4001/003.024.0267 |
| Eriosoma | lanigerum | South Africa | Southern | Hemiptera | 2017 | https://journals.co.za/doi/abs/10.4001/003.025.0123 |
| Frankliniella | occidentalis | South Africa | Southern | Thysanoptera | 2019 | https://hdl.handle.net/10520/EJC-18640d2c5a |

| | | | | | | |
|---|---|---|---|---|---|---|
| Frankliniella | occidentalis | South Africa | Southern | Thysanoptera | 2019 | https://journals.co.za/doi/abs/10.4001/003.027.0322 |
| Holocacista | capensis | South Africa | Southern | Lepidoptera | 2019 | http://www.scielo.org.za/scielo.php?script=sci_arttext&pid=S2224-79042019000200016 |
| Macrotermes | bellicosus | Benin | Western | Isoptera | 2014 | https://brill.com/view/journals/nemy/16/6/article-p719_8.xml |
| Macrotermes | bellicosus | Benin | Western | Isoptera | 2014 | https://brill.com/view/journals/nemy/16/1/article-p109_11.xml |
| Maliarpha | separatella | Kenya | Eastern | Lepidoptera | 2020 | https://www.academia.edu/download/64480239/UJAR3-10414850.pdf |
| Maliarpha | separatella | Kenya | Eastern | Lepidoptera | 2013 | https://www.cabdirect.org/cabdirect/abstract/20133240630 |
| Octodonta | nipae | Nigeria | Western | Coleoptera | 2020 | http://www.esnjournal.com.ng/download/Vol_36/paper_10.pdf |
| Phlyctinus | callosus | South Africa | Southern | Coleoptera | 2019 | https://doi.org/10.1111/aen.12386 |
| Phlyctinus | callosus | South Africa | Southern | Coleoptera | 2013 | https://www.cambridge.org/core/journals/journal-of-helminthology/article/abs/potential-of-[illegible] curculionidae/C8C919A6FEB452786FF633A82A2594CF |
| Phlyctinus | callosus | South Africa | Southern | Coleoptera | 2010 | https://www.cabdirect.org/cabdirect/abstract/20103287864 |
| PHLYCTINUS | callosus | South Africa | Southern | Coleoptera | 2010 | https://www.cabdirect.org/cabdirect/abstract/20103287864 |
| Plangia | ficus | South Africa | Southern | Hemiptera | 2018 | http://dx.doi.org/10.21548/39-2-3158 |
| Planococcus | ficus | South Africa | Southern | Hemiptera | 2015 | http://www.scielo.org.za/pdf/sajev/v36n1/12.pdf |
| Planococcus | ficus | South Africa | Southern | Hemiptera | 2013 | https://doi.org/10.21548/34-1-1086 |
| Planococcus | citri | South Africa | Southern | Hemiptera | 2012 | https://doi.org/10.1016/j.jip.2012.07.023 |
| Planococcus | ficus | South Africa | Southern | Hemiptera | 2013 | https://scholar.archive.org/work/sbbqk3derrbojkc2jaidgjucji/access/wayback/https://www.[illegible] |
| Planococcus | citri | South Africa | Southern | Hemiptera | 2012 | http://scholar.sun.ac.za/handle/10019.1/20147 |
| PLANOCOCCUS | VIBURNI | South Africa | Southern | Hemiptera | 2009 | http://scholar.sun.ac.za/handle/10019.1/2463 |

| | | | | | | |
|---|---|---|---|---|---|---|
| PLANOCOCCUS | citri | South Africa | Southern | Hemiptera | 2015 | https://doi.org/10.1017/S0022149X13000771 |
| Planococcus | xylostella | Benin | Western | Lepidoptera | 2019 | https://dx.doi.org/10.25518/1780-4507.18134 |
| Plutella | xylostella | Ethiopia | Eastern | Lepidoptera | 2016 | https://www.thaiscience.info/Journals/Article/IJAT/10985271.pdf |
| Plutella | xylostella | Kenya | Eastern | Lepidoptera | 2008 | https://doi.org/10.1080/09670870802419636 |
| Plutella | xylostella | Kenya | Eastern | Lepidoptera | 2007 | https://www.cabdirect.org/cabdirect/abstract/20103268557 |
| Plutella | viburni | South Africa | Southern | Hemiptera | 2015 | https://doi.org/10.1080/09670874.2015.1122250 |
| Pseudococcus | ferrugineus | Nigeria | Western | Coleoptera | 2021 | https://ejaj.journals.ekb.eg/article_195928.html |
| Pseudococcus | calamistis | Nigeria | Western | Lepidoptera | 2021 | https://ejaj.journals.ekb.eg/article_195928.html |
| Rhynchophorus | calamistis | Nigeria | Western | Lepidoptera | 2018 | https://www.academia.edu/download/57019694/G1106024853.pdf |
| Sesamia | frugiperda | Ghana | Western | Lepidoptera | 2021 | https://dx.doi.org/10.21608/ejaj.2021.192214 |
| Sesamia | frugiperda | Nigeria | Western | Lepidoptera | 2021 | https://ejaj.journals.ekb.eg/article_195928.html |
| Spodoptera | frugiperda | Rwanda | Eastern | Lepidoptera | 2020 | https://www.researchgate.net/profile/Stefan-Toepfer/publication/341553716_A_Rwandan_survey_of_entomopathogenic_nematodes survey-of-entomopathogenic-nematodes-that-can-potentially-be-used-to-control-the-fall-a |
| Spodoptera | vayssierei | Cameroon | Central | Hemiptera | 2020 | https://doi.org/10.1016/j.biocontrol.2020.104321 |
| Spodoptera | leucotreta | South Africa | Southern | Lepidoptera | 2011 | https://doi.org/10.1016/j.jip.2011.07.006 |
| Spodoptera | leucotreta | South Africa | Southern | Lepidoptera | 2018 | https://journals.co.za/doi/abs/10.4001/003.026.0014 |
| Spodoptera | leucotreta | South Africa | Southern | Lepidoptera | 2016 | https://journals.co.za/doi/abs/10.4001/003.024.0489 |
| Stictococcus | leucotreta | South Africa | Southern | Lepidoptera | 2017 | https://doi.org/10.1080/09583157.2017.1391174 |
| Thaumatotibia | leucotreta | South Africa | Southern | Lepidoptera | 2013 | https://doi.org/10.1080/09583157.2013.854316 |
| Thaumatotibia | leucotreta | South Africa | Southern | Lepidoptera | 2019 | https://doi.org/10.1007/s10526-019-09943-3 |
| Thaumatotibia | occidentalis | Benin | Western | Isoptera | 2014 | https://brill.com/view/journals/nemy/16/6/article-p719_8.xmll |

| | | | | | | |
|---|---|---|---|---|---|---|
| Thaumatotibia | absoluta | Kenya | Eastern | Lepidoptera | 2017 | https://www.academia.edu/download/56118768/wjar-5-4-5.pdf |
| Thaumatotibia | absoluta | Kenya | Eastern | Lepidoptera | 2018 | https://www.researchgate.net/profile/David-Munyua-2/publication/322963834_Integrated_use_of_Kenyan_Entomopathogenic_Nematodes_S use-of-Kenyan-Entomopathogenic-Nematodes-Steinernema-Species-and-Neem-Against |
| Thaumatotibia | absoluta | Kenya | Eastern | Lepidoptera | 2021 | https://scholar.archive.org/work/f437ga5asfdv3lty26h5mrdj4u/access/wayback/https://ccs |
| Trinervitermes | absoluta | Rwanda | Eastern | Lepidoptera | 2019 | https://doi.org/10.1186/s41938-019-0163-3 |
| Tuta | absoluta | Rwanda | Eastern | Lepidoptera | 2020 | https://doi.org/10.1016/j.cropro.2020.105183 |
| Tuta | absoluta | South Africa | Southern | Lepidoptera | 2020 | http://repository.nwu.ac.za/bitstream/handle/10394/36586/Coleman_O.pdf?sequence=1& |
| Tuta | *grubs* | Rwanda | Eastern | Coleoptera | 2018 | https://doi.org/10.1186/s41938-017-0003-2 |
| Tuta | grubs | South Africa | Southern | Coleoptera | 2019 | https://doi.org/10.1016/j.biocontrol.2019.104043 |
| Tuta | frugiperda | Ghana | Western | Lepidoptera | 2023 | https://www.ajol.info/index.php/acsj/article/view/236793 |
| Tuta | frugiperda | Rwanda | Eastern | Lepidoptera | 2022 | https://www.mdpi.com/2075-4450/13/2/205 |
| *Pseudococcus* | viburni | South Africa | Southern | Hemiptera | 2022 | https://doi.org/10.1080/09583157.2021.2010653 |
| Plangia | gramínea | South Africa | Southern | Orthoptera | 2022 | http://dx.doi.org/10.21548/43-2-4897 |

**Table S2: Publications on EPN- related research in SSA, year 2003-2022**

| Country | Region | year | DOI / Source link |
|---|---|---|---|
| Benin | Western | 2004 | https://brill.com/view/journals/nemy/17/9/article-p1057_5.xml |
| Benin | Western | 2005 | https://doi.org/10.1016/j.biocontrol.2017.10.009 |

| | | | |
|---|---|---|---|
| Benin | Western | 2006 | https://www.researchgate.net/profile/Lieven-Waeyenberge/publication/259531677_First_record_on_the_distribution_of_entomopathog on-the-distribution-of-entomopathogenic-nematodes-Rhabditida-Steinernematidae-and-Heterorhabditidae-in-Southern-Benin.pdff |
| Benin | Western | 2006 | https://brill.com/view/journals/nemy/21/2/article-p107_1.xml |
| Benin | Western | 2007 | https://doi.org/10.1007/s10526-014-9568-9 |
| Benin | Western | 2008 | https://brill.com/view/journals/nemy/16/6/article-p719_8.xml |
| Benin | Western | 2008 | https://www.entomoljournal.com/archives/2017/vol5issue6/PartH/5-4-307-961.pdf |
| Benin | Western | 2008 | https://www.ajol.info/index.php/aga/article/view/111255 |
| Benin | Western | 2008 | https://dx.doi.org/10.25518/1780-4507.18134 |
| Benin | Western | 2009 | https://doi.org/10.1007/s00203-017-1470-2 |
| Benin | Western | 2009 | https://doi.org/10.1016/j.cropro.2021.105754 |
| Benin | Western | 2009 | https://doi.org/10.4314/ijbcs.v13i1.36 |
| Benin | Western | 2009 | https://doi.org/10.21307%2Fjofnem-2019-066 |
| Benin | Western | 2009 | https://doi.org/10.1186/s41938-021-00448-9 |
| Benin | Western | 2009 | https://brill.com/view/journals/nemy/16/1/article-p109_11.xml |
| Ghana | Western | 2010 | https://dx.doi.org/10.21608/ejaj.2021.192214 |
| Ghana | Western | 2010 | https://www.researchgate.net/profile/Yaw-Danso/publication/342927427_Survey_of_Entomopathogenic_Nematodes_Potential_Biological_Control_Agents_in_Ghanaian_Soils_S of-Entomopathogenic-Nematodes-Potential-Biological-Control-Agents-in-Ghanaian-Soils-Survey-of-Entomopathogenic-Nematodes-Po |
| Nigeria | Western | 2011 | https://journals.flvc.org/nemamedi/article/view/87087 |
| Nigeria | Western | 2011 | https://doi.org/10.1016/S2095-3119(21)63609-2 |
| Nigeria | Western | 2011 | http://icidr.org/jeiadc-vol7no3-dec2015/Isolation%20of%20Entomopathogenic%20Nematode-Bacteria%20Complex%20with%20a%20F |
| Nigeria | Western | 2011 | https://www.researchgate.net/profile/Mohammed-Rufai-2/publication/343037135_Occurrence_of_entomopathogenic_nematodes_in_O Southwestern-Nigeria.pdf |
| Nigeria | Western | 2011 | https://ejaj.journals.ekb.eg/article_195928.html |
| Nigeria | Western | 2011 | https://www.academia.edu/download/57019694/G1106024853.pdf |
| Nigeria | Western | 2012 | https://www.ajol.info/index.php/tjs/article/view/197057/185925 |

| | | | |
|---|---|---|---|
| Nigeria | Western | 2012 | https://www.ajol.info/index.php/jasem/article/view/217962/205561 |
| Nigeria | Western | 2012 | https://www.academia.edu/download/37455562/Nematode_Diversity_on_some_common_weeds_in_Makurdi.pdf |
| Nigeria | Western | 2012 | http://www.esnjournal.com.ng/download/Vol_36/paper_10.pdf |
| Cameroon | Central | 2012 | https://doi.org/10.1016/j.jip.2011.09.008 |
| Cameroon | Central | 2013 | https://doi.org/10.1016/j.biocontrol.2020.104321 |
| Ethopia | Eastern | 2013 | https://www.cabdirect.org/cabdirect/abstract/20063062524 |
| Ethopia | Eastern | 2013 | https://brill.com/view/journals/nemy/17/7/article-p741_1.xml |
| Ethopia | Eastern | 2013 | https://brill.com/view/journals/nemy/14/6/article-p741_6.xml |
| Ethopia | Eastern | 2013 | https://www.cabdirect.org/cabdirect/abstract/20113280366 |
| Ethopia | Eastern | 2013 | https://brill.com/view/journals/nemy/6/6/article-p839_5.xml |
| Ethopia | Eastern | 2013 | https://www.thaiscience.info/Journals/Article/IJAT/10985271.pdf |
| Tanzania | Eastern | 2013 | https://doi.org/10.1017/S1742758411000294 |
| Tanzania | Eastern | 2013 | https://doi.org/10.1017/S0022149X15001157 |
| Kenya | Eastern | 2014 | https://doi.org/10.1111/j.1365-2028.2008.00933.x |
| Kenya | Eastern | 2014 | https://doi.org/10.1080/09670870802419636 |
| Kenya | Eastern | 2014 | https://www.cabdirect.org/cabdirect/abstract/20093232109 |
| Kenya | Eastern | 2014 | https://www.academia.edu/download/56118768/wjar-5-4-5.pdf |
| Kenya | Eastern | 2014 | https://www.cabdirect.org/cabdirect/abstract/20103268557 |
| Kenya | Eastern | 2014 | https://www.cabdirect.org/cabdirect/abstract/20163288738 |
| Kenya | Eastern | 2014 | https://www.researchgate.net/profile/David-Munyua-2/publication/322963834_Integrated_use_of_Kenyan_Entomopathogenic_Nemato<br>Entomopathogenic-Nematodes-Steinernema-Species-and-Neem-Against-Tuta-Absoluta-on-Tomato.pdf |
| Kenya | Eastern | 2014 | https://pdfs.semanticscholar.org/d108/71f837102af8b13b1d798b3fb6b27c15a559.pdf |
| Kenya | Eastern | 2014 | https://scholar.archive.org/work/f437ga5asfdv3lty26h5mrdj4u/access/wayback/https://ccsenet.org/journal/index.php/jas/article/downloa |
| Kenya | Eastern | 2014 | https://ideas.repec.org/a/aoj/agafsr/v3y2016i1p29-36id160.html |

| | | | |
|---|---|---|---|
| Kenya | Eastern | 2014 | https://www.ajol.info/index.php/jagst/article/view/219191/206835 |
| Kenya | Eastern | 2014 | https://www.kalro.org/www.eaafj.or.ke/index.php/path/article/view/390 |
| Kenya | Eastern | 2014 | https://www.academia.edu/download/64480239/UJAR3-10414850.pdf |
| Kenya | Eastern | 2015 | https://www.cabdirect.org/cabdirect/abstract/20133240630 |
| Kenya | Eastern | 2015 | http://ir-library.mmust.ac.ke:8080/jspui/bitstream/190/278/1/Kawaka%20Fanuel.pdf |
| Kenya | Eastern | 2015 | http://www.scielo.org.mx/scielo.php?pid=S1870-04622011000400010&script=sci_arttext&tlng=en |
| Kenya | Eastern | 2015 | http://erepository.uonbi.ac.ke/handle/11295/26308 |
| Kenya | Eastern | 2015 | https://academicjournals.org/journal/AJAR/article-full-text-pdf/543EADD36101 |
| Kenya | Eastern | 2015 | http://erepository.uonbi.ac.ke/handle/11295/14404 |
| Kenya | Eastern | 2015 | http://erepository.uonbi.ac.ke/handle/11295/104268 |
| Rwandan | Eastern | 2015 | https://doi.org/10.1186/s41938-017-0003-2 |
| Rwandan | Eastern | 2015 | https://doi.org/10.1080/09583157.2016.1159658 |
| Rwandan | Eastern | 2015 | https://doi.org/10.1186/s41938-019-0163-3 |
| Rwandan | Eastern | 2015 | https://www.researchgate.net/profile/Stefan-Toepfer/publication/341553716_A_Rwandan_survey_of_entomopathogenic_nematodes_th nematodes-that-can-potentially-be-used-to-control-the-fall-armyworm.pdf |
| Rwandan | Eastern | 2015 | https://sciendo.com/it/article/10.21307/jofnem-2021-089 |
| Rwandan | Eastern | 2015 | https://doi.org/10.1016/j.cropro.2020.105183 |
| South Africa | Southern | 2016 | https://journals.co.za/doi/epdf/10.10520/EJC87795 |
| South Africa | Southern | 2016 | https://doi.org/10.1016/j.jip.2011.07.006 |
| South Africa | Southern | 2016 | https://doi.org/10.1016/j.jip.2009.07.003 |
| South Africa | Southern | 2016 | https://doi.org/10.1007/978-3-319-18266-7_20 |
| South Africa | Southern | 2016 | https://scholar.sun.ac.za/bitstream/10019.1/85898/1/malan_heterorhabditis_2013.pdf |

| | | | |
|---|---|---|---|
| South Africa | Southern | 2016 | https://brill.com/view/journals/nemy/8/2/article-p157_1.xml |
| South Africa | Southern | 2016 | http://www.scielo.org.za/pdf/sajev/v41n1/01.pdf |
| South Africa | Southern | 2016 | https://www.cambridge.org/core/journals/journal-of-helminthology/article/abs/steinernema-jeffreyense-n-sp-rhabditida-steinernematidae |
| South Africa | Southern | 2016 | https://brill.com/view/journals/nemy/10/3/article-p381_7.xml |
| South Africa | Southern | 2016 | https://www.researchgate.net/profile/S-Patricia-Stock/publication/263711042_Steinernema_tophus_sp_n_Nematoda_Steinernematidae Steinernematidae-a-new-entomopathogenic-nematode-from-South-Africa.pdf |
| South Africa | Southern | 2016 | https://journals.co.za/doi/epdf/10.10520/EJC150979 |
| South Africa | Southern | 2016 | https://brill.com/view/journals/nemy/13/5/article-p569_7.xml |
| South Africa | Southern | 2016 | https://journals.co.za/doi/abs/10.4001/003.026.0014 |
| South Africa | Southern | 2016 | https://brill.com/view/journals/nemy/22/3/article-p343_7.xml |
| South Africa | Southern | 2016 | https://www.cambridge.org/core/journals/journal-of-helminthology/article/steinernema-innovationi-n-sp-panagrolaimomorpha-steinerner |
| South Africa | Southern | 2017 | https://brill.com/view/journals/nemy/16/4/article-p475_10.xml |
| South Africa | Southern | 2017 | https://brill.com/view/journals/nemy/18/4/article-p439_4.xml |
| South Africa | Southern | 2017 | https://journals.co.za/doi/abs/10.4001/003.024.0489 |
| South Africa | Southern | 2017 | https://sciendo.com/es/article/10.21307/jofnem-2017-022 |
| South Africa | Southern | 2017 | https://www.tandfonline.com/doi/abs/10.1080/03235408.2018.1475281 |
| South Africa | Southern | 2017 | https://www.tandfonline.com/doi/abs/10.1080/03235408.2021.1931648 |
| South Africa | Southern | 2017 | https://brill.com/view/journals/nemy/18/5/article-p571_5.xml |

| | | | |
|---|---|---|---|
| South Africa | Southern | 2018 | https://www.researchgate.net/profile/Mahloro-Serepa-Dlamini/publication/342374998_Molecular_identification_of_a_Heterorhabditis_e identification-of-a-Heterorhabditis-entomopathogenic-nematode-isolated-from-the-northernmost-region-of-South-Africa.pdf |
| South Africa | Southern | 2018 | https://brill.com/view/journals/nemy/20/4/article-p355_3.xml |
| South Africa | Southern | 2018 | https://www.tandfonline.com/doi/abs/10.1080/03235408.2019.1588193 |
| South Africa | Southern | 2018 | https://journals.co.za/doi/abs/10.10520/EJC155713 |
| South Africa | Southern | 2018 | https://doi.org/10.1080/09583157.2017.1391174 |
| South Africa | Southern | 2018 | https://brill.com/view/journals/nemy/18/2/article-p235_9.xml |
| South Africa | Southern | 2018 | https://journals.co.za/doi/abs/10.10520/EJC176592 |
| South Africa | Southern | 2018 | https://doi.org/10.1016/j.cropro.2017.11.008 |
| South Africa | Southern | 2018 | https://doi.org/10.1016/j.biocontrol.2019.104043 |
| South Africa | Southern | 2018 | https://journals.co.za/doi/abs/10.4001/003.024.0061 |
| South Africa | Southern | 2018 | https://brill.com/view/journals/nemy/19/10/article-p1157_4.xml |
| South Africa | Southern | 2018 | https://hdl.handle.net/10520/EJC195098 |
| South Africa | Southern | 2018 | http://dx.doi.org/10.17159/sajs.2019/6008 |
| South Africa | Southern | 2019 | https://doi.org/10.1111/aen.12386 |
| South Africa | Southern | 2019 | https://hdl.handle.net/10520/EJC-18640d2c5a |
| South Africa | Southern | 2019 | http://dx.doi.org/10.21548/39-2-3158 |
| South Africa | Southern | 2019 | https://journals.co.za/doi/epdf/10.10520/EJC176580 |

| | | | |
|---|---|---|---|
| South Africa | Southern | 2019 | https://doi.org/10.1016/j.gdata.2016.01.017 |
| South Africa | Southern | 2019 | http://www.scielo.org.za/pdf/sajev/v36n1/12.pdf |
| South Africa | Southern | 2019 | https://www.cambridge.org/core/journals/journal-of-helminthology/article/abs/potential-of-entomopathogenic-nematodes-for-the-control- |
| South Africa | Southern | 2019 | https://doi.org/10.1371/journal.pone.0242645 |
| South Africa | Southern | 2019 | https://doi.org/10.21548/34-1-1086 |
| South Africa | Southern | 2019 | https://doi.org/10.1099/ijs.0.049049-0 |
| South Africa | Southern | 2019 | https://doi.org/10.1007/s10526-019-09945-1 |
| South Africa | Southern | 2019 | https://www.cambridge.org/core/journals/journal-of-helminthology/article/abs/entomopathogenic-nematodes-for-the-control-of-the-codli |
| South Africa | Southern | 2019 | https://doi.org/10.1016/j.jip.2012.07.023 |
| South Africa | Southern | 2019 | https://academicjournals.org/journal/JEN/article-full-text-pdf/243CF0B60140 |
| South Africa | Southern | 2019 | https://journals.co.za/doi/epdf/10.4001/003.026.0337 |
| South Africa | Southern | 2019 | https://scholar.archive.org/work/sbbqk3derrbojkc2jaidgjucji/access/wayback/https://www.journals.ac.za/index.php/sajev/article/downloa |
| South Africa | Southern | 2019 | https://journals.co.za/doi/abs/10.4001/003.025.0123 |
| South Africa | Southern | 2020 | https://www.researchgate.net/profile/Leon-Dicks/publication/264746171_First_report_of_the_symbiotic_bacterium_Xenorhabdus_indica symbiotic-bacterium-Xenorhabdus-indica-associated-with-the-entomopathogenic-nematode-Steinernema-yirgalemense.pdf |
| South Africa | Southern | 2020 | https://journals.co.za/doi/abs/10.4001/003.027.0322 |
| South Africa | Southern | 2020 | https://journals.co.za/doi/abs/10.10520/EJC155697 |
| South Africa | Southern | 2020 | https://journals.co.za/doi/abs/10.10520/EJC150975 |

| | | | |
|---|---|---|---|
| South Africa | Southern | 2020 | http://www.scielo.org.za/scielo.php?script=sci_arttext&pid=S2224-79042019000200016 |
| South Africa | Southern | 2020 | https://doi.org/10.1080/09583157.2013.854316 |
| South Africa | Southern | 2020 | https://doi.org/10.1080/09583157.2016.1217393 |
| South Africa | Southern | 2020 | https://doi.org/10.1080/09670874.2015.1122250 |
| South Africa | Southern | 2020 | https://doi.org/10.3390/insects10040117 |
| South Africa | Southern | 2020 | https://peerj.com/articles/1023/ |
| South Africa | Southern | 2020 | http://scholar.sun.ac.za/handle/10019.1/20147 |
| South Africa | Southern | 2020 | http://www.scielo.org.za/pdf/sajev/v42n2/03.pdf |
| South Africa | Southern | 2021 | http://scholar.sun.ac.za/handle/10019.1/2463 |
| South Africa | Southern | 2021 | https://doi.org/10.1017/S0022149X13000771 |
| South Africa | Southern | 2021 | https://doi.org/10.1007/s10526-019-09943-3 |
| South Africa | Southern | 2021 | https://doi.org/10.1080/09583157.2011.607922 |
| South Africa | Southern | 2021 | http://www.scielo.org.za/pdf/sajev/v41n2/06.pdf |
| South Africa | Southern | 2021 | https://doi.org/10.1080/03235408.2021.1914369 |
| South Africa | Southern | 2021 | https://www.proquest.com/openview/04135b98c2c95a02ccc71927f764795d/1?pq-origsite=gscholar&cbl=886351 |
| South Africa | Southern | 2021 | https://journals.co.za/doi/abs/10.4001/003.026.0001 |
| South Africa | Southern | 2021 | https://doi.org/10.1080/03235408.2021.1899357 |

| | | | |
|---|---|---|---|
| South Africa | Southern | 2021 | https://journals.tubitak.gov.tr/cgi/viewcontent.cgi?article=1062&context=zoology |
| South Africa | Southern | 2021 | https://doi.org/10.1016/j.gene.2021.145780 |
| South Africa | Southern | 2021 | https://sciendo.com/pdf/10.21307/jofnem-2018-049 |
| South Africa | Southern | 2021 | https://koedoe.co.za/index.php/koedoe/article/download/1661/2892 |
| South Africa | Southern | 2021 | https://doi.org/10.1016/j.jip.2008.09.007 |
| South Africa | Southern | 2021 | http://repository.nwu.ac.za/bitstream/handle/10394/36586/Coleman_O.pdf?sequence=1&isAllowed=y |
| South Africa | Southern | 2021 | https://www.cabdirect.org/cabdirect/abstract/20103287864 |
| South Africa | Southern | 2021 | http://scholar.sun.ac.za/handle/10019.1/4113 |
| Ghana | Western | 2022 | https://www.ajol.info/index.php/acsj/article/view/236793 |
| Rwandan | Eastern | 2022 | https://sciendo.com/es/article/10.2478/jofnem-2022-0049 |
| Rwandan | Eastern | 2022 | https://www.mdpi.com/2075-4450/13/2/205 |
| South Africa | Southern | 2022 | https://doi.org/10.1080/09583157.2022.2099528 |
| South Africa | Southern | 2022 | https://www.cambridge.org/core/services/aop-cambridge-core/content/view/0A3835E1CB560A818E5DE949B3DD3F28/S0022149X22[illegible] |
| South Africa | Southern | 2022 | https://doi.org/10.1080/09583157.2021.2010653 |
| South Africa | Southern | 2022 | http://dx.doi.org/10.21548/43-2-4897 |
